# The Effect of Intracerebroventricular Fibroblast Growth Factor 23 on gene expression in the Rats’ Hypothalamus

**DOI:** 10.1101/2022.11.18.516150

**Authors:** Stan R Ursem, Charlene Diepenbroek, Tess Kool, Leslie Eggels, Annemieke C Heijboer, Susanne E la Fleur

## Abstract

Fibroblast growth factor 23 (FGF23) is a key regulator of systemic phosphate homeostasis, but also an interplay with glucose metabolism has been suggested. Several studies implicate a function of FGF23 in the brain, and indeed we have recently identified FGF23 protein in several brain areas in rats, such as the hypothalamus, third ventricle and choroid plexus. In the current study, we aimed to determine the effect of an intracerebroventricular (icv) injection of FGF23 in the third ventricle of rats on hypothalamic genes involved in glucose regulation. In addition, we assessed whether glycerol can be used safely for icv injections as glycerol is used as a stabilizing compound for FGF23 protein.

Adult Wistar rats received an icv injection of recombinant rat FGF23 or vehicle. Dose dependent behavioral changes, suggestive of stress, were observed directly after infusion of FGF23. After 60 min animals were sacrificed and the arcuate nucleus, lateral hypothalamus and choroid plexus were isolated. In these brain regions gene expression was determined of the FGF23 receptor complex (FGFR1, αKlotho), NPY, POMC, phosphate transporters (SLC20 and SLC34 families) and markers of cellular ER stress (ATF4 and the ratio of spliced/unspliced XBP1).

We showed that glycerol is well tolerated as stabilizer for icv injections. In FGF23-treated animals, cellular ER stress markers were increased in the arcuate nucleus. FGF23 injection did not affect expression of its receptor complex, NPY, POMC, or phosphate transporters. Future studies are warranted to investigate the effect of FGF23 in the brain on the protein level and on neuronal activation.

## 1. Introduction

Fibroblast growth factor 23 (FGF23) is an endocrine FGF, best known for its phosphaturic effects.^1^ It is produced primarily by osteocytes in the bone and in order to exert its phosphate regulating role, FGF23 binds to the FGF receptor 1 (FGFR1) in the kidney. FGF23 itself has a low affinity for the receptor and needs presence of the co-receptor αKlotho to induce downstream signaling.

Next to its classical function – renal regulation of serum phosphate – FGF23 secretion was found to be altered by glucose and insulin, suggesting an interplay with glucose metabolism.^2,3^ In a recent experiment in humans, an oral glucose tolerance test altered FGF23 concentrations, which was not explained by a change in serum phosphate.^4^ This is in line with observational studies pointing towards a link between FGF23 and insulin resistance.^5–8^ Subsequent studies confirmed the interplay between insulin, glucose and FGF23 *in vitro*, which further suggests a possible relation between FGF23 and glucose homeostasis.^2,3^

An important regulator of glucose homeostasis is the brain. Interestingly, the needed receptor complex for classical FGF23 signaling, being FGFR1 and αKlotho, has been shown to be present throughout the brain.^9,10^ Thus, it could well be that the brain is a target of extra-renal FGF23 action.^11–13^ In fact, the study which first identified FGF23 gene expression, reported localization in the ventrolateral thalamic nucleus.^14^ Unfortunately, subsequent studies were inconsistent in reporting the presence of FGF23 mRNA in the brain. Therefore, we recently localized and visualized FGF23 expression in the brain with focus on the hypothalamus, being an important area for glucose regulation. Using a specific monoclonal FGF23 antibody, we observed FGF23 protein in the hypothalamus and more specifically in the arcuate nucleus, as well as in the third ventricle lining and choroid plexus of rats.^15^ However, no FGF23 gene expression was observed, suggesting the detected protein in the brain is from peripheral origin and is thus probably bone-derived. In humans, FGF23 protein has been found in CSF.^16,17^ With the confirmation of the protein being present in the brain, exciting questions arise on its potential function.

A growing body of evidence shows an important relation between fibroblast growth factors, the brain and glucose metabolism.^18,19^ Central injection of FGF1 in the ventricle of rodents was found to induce a sustained remission of a diabetic phenotype.^20^ Tanycytes, specialized cells located in the lower part of the third ventricle, appeared to play a key role in the underlying mechanism. Besides FGF23, there are two other endocrine FGFs: FGF19 (of which FGF15 is the rodent orthologue) and FGF21. These were also found to have potential therapeutic applications for type 2 Diabetes, which are in part brain-mediated through the hypothalamus.^21^ In light of these promising reports, it is intriguing that preliminary data indeed suggest a central effect of FGF23 on two neuronal populations located in the hypothalamus.^22^

In the hypothalamus, and more specifically in the arcuate nucleus, nutrients and hormonal signals related to energy metabolism can easily cross the blood brain barrier and affect different neuronal populations. Activation of these by nutrients and several humoral factors regulates feeding behavior, energy expenditure and thermoregulation, but these neurons have also been shown to change glucose production and insulin sensitivity.^23,24^ Within the arcuate nucleus two neuronal populations are of special physiological importance: the orexigenic Neuropeptide Y (NPY) and Agouti Related Peptide (AgRP) expressing neurons and the anorexigenic Proopiomelanocortin (POMC) expressing neurons. Central administration of NPY has been shown to reduce insulin sensitivity, whereas melanocortin stimulating hormone enhances the action of insulin on glucose uptake and production.^25,26^ Interestingly, a published abstract showed that intracerebroventricular (icv) injection of FGF23 in mice resulted in increased gene expression of NPY and POMC in the hypothalamus.^22^ Thus, we hypothesize that FGF23 can regulate glucose metabolism via NPY and POMC neurons in the arcuate nucleus.

Our study objective was to examine the emerging role of FGF23 in the brain, and its role in glucose metabolism. Therefore, we first investigated in male rats the effect of an icv FGF23 injection on expression of genes involved in glucose regulation in the arcuate nucleus, being NPY and POMC to verify the preliminary data reported in mice. Moreover, based on our previous results on FGF23 protein expression in the brain, also the lateral hypothalamus and the choroid plexus were included in analysis to determine whether the components of the receptor complex respond to FGF23 infusion. In addition, phosphate transporter gene expression was included, as this is an important function of FGF23 in the periphery. During the experiment a behavioral response was observed upon FGF23 infusion, because of which also cellular stress markers were included in the study.

## 2. Methods

### 2.1. Animals

Male adult Wistar rats (Charles River, Germany), weighing 260-320 g were used. Rats were housed in the animal facility of the Netherlands Institute of Neuroscience in a temperature and light-controlled room (21-23°C; 12:12h light/dark cycle, 07:00-19:00 lights on). *Ad libitum* access to a standard diet (Teklad global diet 2918; 24% protein, 58% carbohydrate, and 18% fat, 3.1 kcal/g, Envigo) and a bottle of tap water was given. The animal care committee of the Netherlands Institute for Neuroscience approved all experiments according to Dutch legal ethical guidelines.

### 2.2. Surgery

One week after arrival of the rats, a cannula was implanted in the third ventricle of the brain. Rats were anesthetized with a mixture of intraperitoneal ketamine (80 mg/kg; Eurovet Animal Health, Blader, The Netherlands), xylazine (8 mg/kg; Bayer Health Care, Mijdrecht, The Netherlands) and atropine (0.1 mg/kg; Pharmacie B.V. Haarlem, The Netherlands). Subsequently, rats were fixed in a stereotactic frame. A permanent 11 mm long 26-gauge stainless steel cannula (Plastics One, Bilaney Consultants GmbH, Düsseldorf, Germany) was implanted at AP: +0.3 mm, ML: +0.0 mm, DV −9.0 mm (coordinates from Bregma with toothbar setting: 5+, DV coordinates from dura mater). Using four anchor screws and dental cement, guide cannulas were secured to the skull. Guide cannulas were occluded using a 28-gauge stainless steel dummy cannula (Plastics One, Bilaney Consultants GmbH, Dusseldorf, Germany). After surgery and after 24h, rats received an analgesic (carprofen; 0.5 mg/100 g s.c.) and were housed solitarily. Rats recovered for 7 days.

### 2.3. Glycerol injection

The used recombinant FGF23 needed to be dissolved in a 10% glycerol solution, in order to ensure its stability. Therefore, prior to FGF23 injection experiments, in a separate group of rats it was studied whether a 10% glycerol solution was safe for icv injection and did not exert effects on body weight, water consumption or food intake. Six rats received an infusion of 1.5 μL (n = 4) or 3.0 μL (n = 2) of 10% glycerol (Merck, Darmstadt, Germany) in phosphate-buffered saline (PBS) or PBS alone. Every animal received two injections of either glycerol or vehicle (PBS) in random order, serving as their own control, with an interval of one week.

Injections were given at the same time point in the morning (two hours after lights on), animals were fasted overnight. Injectors extended 1 mm below the end of the cannula and were connected to a 10 μL Hamilton syringe. Using an infusion pump (Harvard Apparatus, Massachusetts, USA) volumes were infused at rate of 0.6 μL/min. Fluid movement was confirmed by a small air bubble in the tubing and the injector was left in place for 1 min after infusion to ensure fluid diffusion. After completion, food was returned to the cage.

After 2h, 5h, 24h and 48h rats were monitored and water and food intake were measured. Body weight was measured after 24h and 48h. At the end of the experiment, animals received an icv injection of 1.5 μL Evans-Blue dye with the same settings as described above. Five minutes after infusion, rats were decapitated after 33% CO2 / 66% O2 anesthesia. Brains were rapidly dissected, frozen on dry ice and stored at −80°C. Cannula placement was checked by localizing the Evans-Blue dye in coronal sections of the brain. One animal was excluded from analysis due to cannula misplacement.

### 2.4. FGF23 injection

After placement of third ventricle cannulas and recovery, as detailed above, rats received an injection of FGF23 or vehicle (10% glycerol in PBS). Recombinant rat FGF23 protein (QP6045, enquire Bioreagents, Littleton, USA) was dissolved in PBS with 10% glycerol with pH adjustment to a concentration of 1.0 μg/μL (pH = 8) and a concentration of 0.62 μg/μL (pH = 7.5); the maximal concentration at pH = 7.5 was 0.62 μg/μL. Rats were randomly allocated to receive either FGF23 solution (administered dose and concentration are mentioned in the results section), or a similar volume of vehicle. The doses were estimated considering intravenous infusion of FGF23 in previous studies, the mean basal concentration of intravenous FGF23 compared to the concentration in CSF, and studies administering icv FGF19 or FGF21 in rats.^27–29^

Injections were given in the morning after an overnight fast (two hours after lights on), took place on two separate days. The overnight fast was included as also a fast was used in the earlier published preliminary data.^22^ Using an infusion pump (Harvard Apparatus, Massachusetts, USA) volumes were infused at a rate of 0.6 μL/min, the injector was left in place for 1 min to allow for diffusion. Sixty minutes after removal of the injector, animals were sacrificed by decapitation after 33% CO2 / 66% O2 anesthesia. Brains were rapidly dissected, frozen on dry ice and stored at −80°C for gene expression analysis.

### 2.5. qPCR

Using a cryostat, coronal brain sections were cut (200 μm) and punches were obtained for the arcuate nucleus, lateral hypothalamus and choroid plexus. Punches were obtained with a 1 mm-diameter blunt needle, bilaterally for the lateral hypothalamus and choroid plexus. The following coordinates were used: arcuate nucleus (Bregma −1.68 to −3.28), lateral hypothalamus (Bregma −1.28 to −2.88) and choroid plexus (Bregma −0.48 to −2.08). Sections were put in RNAlater (Ambion, Waltham, USA). After isolation of the punches, these were put on dry ice and stored at −80°C.

Tissue was homogenized in 300 μL TriReagent and total RNA isolated using the Direct-zol RNA Miniprepkit (R2052, Zymo Research, Irvine, USA) with DNAse treatment. The yielded RNA concentration was measured using the Nanodrop (Nanodrop, Wilmington, USA) and RNA quality analyzed on the Bioanalyser (Agilent, Santa Clara, USA). All RIN values were larger than 7. Using the transcriptor first-strand cDNA synthesis kit with oligod(T) primers (04897030001; Roche Molecular Biochemicals, Mannheim, Germany) cDNA was synthesized with equal RNA input (400ng). For several samples no reverse transcriptase was added, to check for genomic DNA contamination.

All qPCRs were performed *in duplo*. Expression of *Atf4*, *αKlotho*, *Fgfr1*, *Slc20a1*, *Slc20a2*, *Slc34a1*, *Slc34a2* and the ratio of spliced/unspliced *Xbp1* was investigated in the lateral hypothalamus and choroid plexus. In addition, *Npy* and *Pomc* expression was assessed in the arcuate nucleus. The following reference genes were used for all areas: *Cyclophilin*, *Hprt*, *β-actin* and *Gapdh*. Primer sequences are mentioned in Table 1. RT-qPCR was performed with 1/10 diluted cDNA with the SensiFAST SYBR no-rox kit (Bioline, London, UK) and Lightcycler^®^ 480 (Roche Molecular Biochemicals, Mannheim, Germany). The PCR product size was visualized on DNA agarose gel.

**Table 1.**
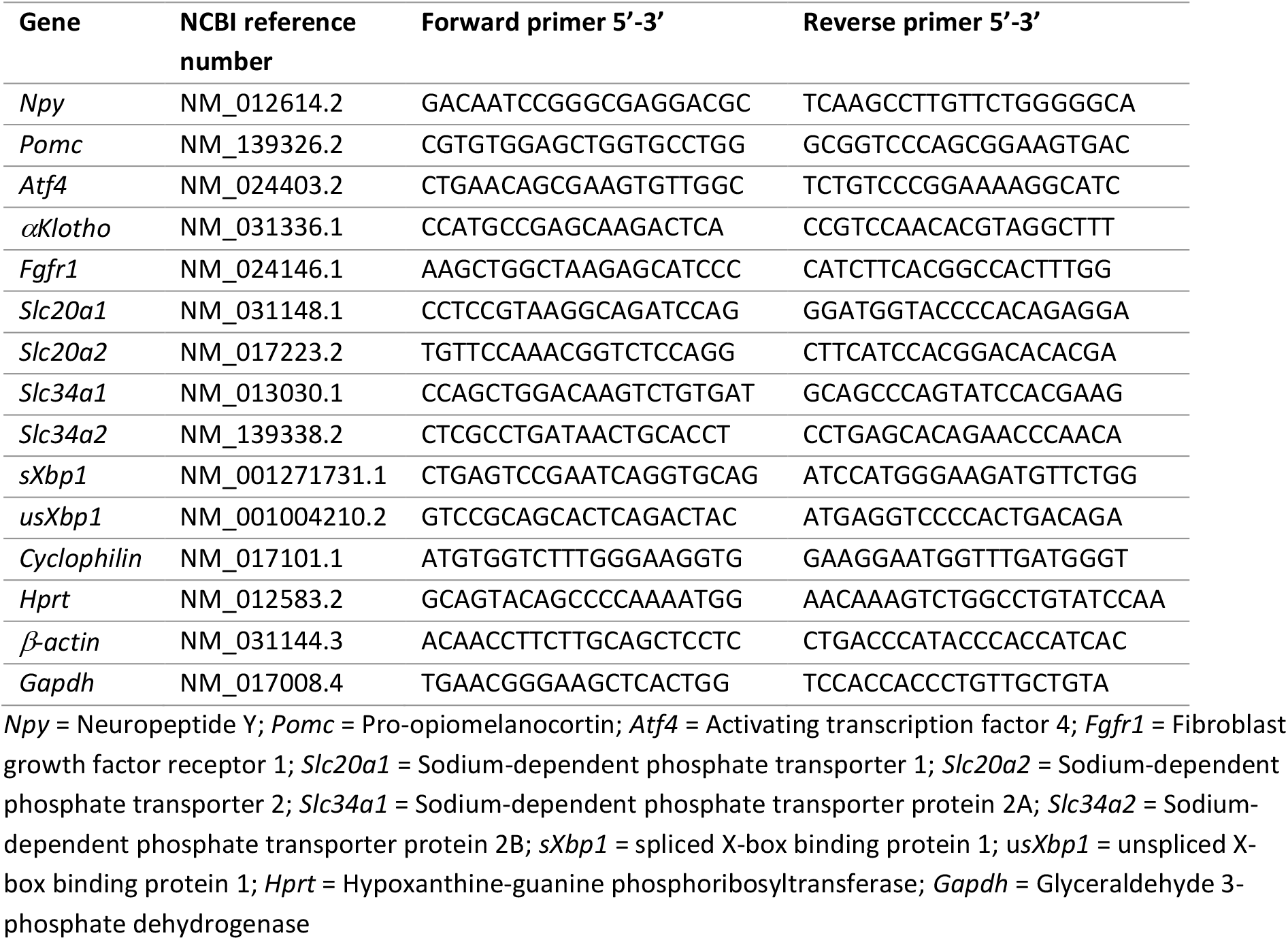
Primer sequences

### 2.6. Statistics and analyses

RT-qPCR quantification was performed using LinReg Software.^30^ Samples having a deviation > 7% from the mean PCR efficiency were excluded. Values were normalized using the geometric mean of the four reference genes.

A paired student’s t-test (glycerol experiment) or Mann-Whitney U test (FGF23 experiment) was used to assess differences. Results are presented as median with interquartile range (IQR) or as mean with standard deviation (SD), dependent on distribution. Statistical analyses were performed using GraphPad Prism 8 (version 8.3.0 [538]). A p-value < 0.05 was considered to reflect statistical significance.

## 3. Results

### 3.1. Effects of icv glycerol injection

During and after infusion of 1.5 μL or 3.0 μL glycerol (10% in PBS) no behavioral changes were observed. For analysis of body weight, food consumption and water intake, the results of 1.5 μL and 3.0 μL were pooled (Figure 1). All interval measurements, shown in Figure 1, exhibited no significant difference between the glycerol or vehicle injected groups. Both groups gained body weight, of which the most within the first 24h, because of reintroduction of chow after an overnight fast. The cumulative effects on body weight, food consumption and water intake also did not differ significantly between the two groups (data not shown). The results indicate that 10% glycerol in PBS is well-tolerated.

**Figure 1.**
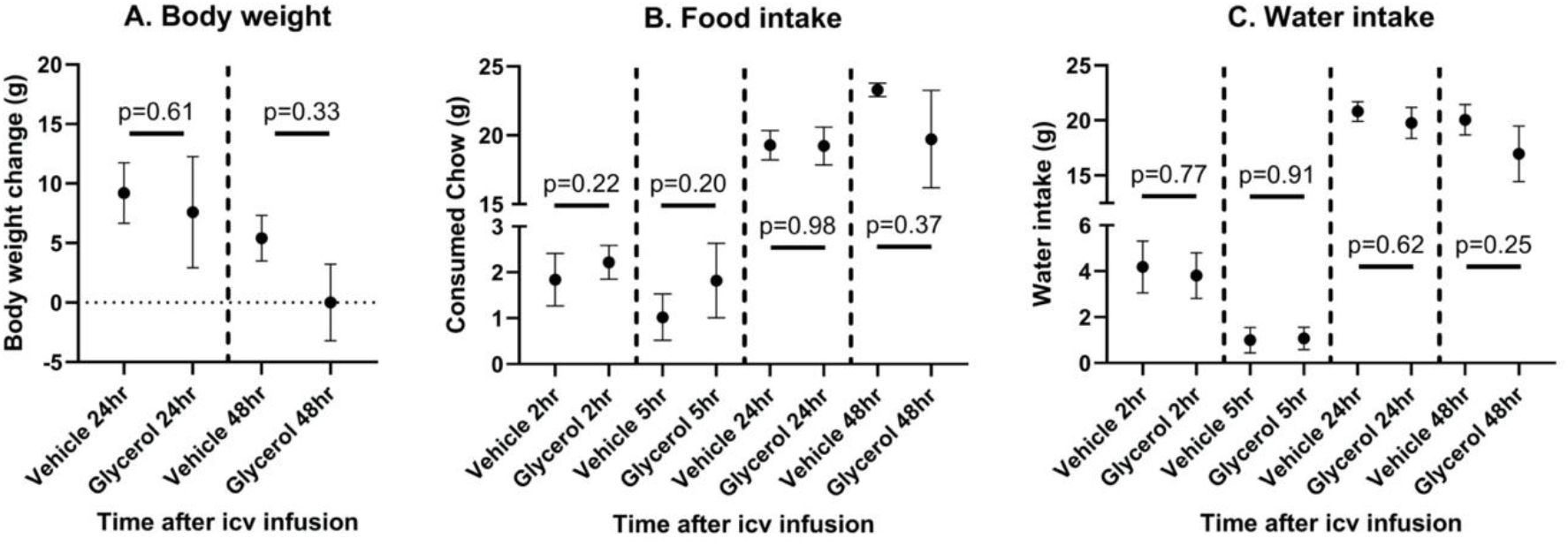
The effect of icv glycerol (10% in PBS) injection on (A) body weight change, (B) food intake and (C) water intake compared to vehicle. Mean values with SEM.

### 3.2. Effects of icv FGF23 injection

The first rat to be infused received 2 μg of recombinant FGF23 (solution 1.00 μg/μL, pH = 8). The infusion of this concentration immediately increased locomotor activity, jumping and escape behavior. Next, we lowered the pH of the solution and infused a lower dose of 1.2 μg of recombinant FGF23 (solution 0.62 μg/μL, pH = 7.5), however some animals still showed similar increased locomotor activity and jumping behavior. Thereafter, the dose was lowered to 0.77 μg (solution 0.62 μg/μL, pH = 7.5), which did not result in behavioral responses. Data from animals not showing behavioral responses (n = 4, dose 1.2 μg or 0.77 μg) were pooled and used for analyzing hypothalamic gene expression.

Gene expression was assessed in the arcuate nucleus, lateral hypothalamus and choroid plexus. The geometric mean of the reference genes did not differ significantly between FGF23 and vehicle treated groups for all three punched regions (data not shown). The normalized mRNA expression for the arcuate nucleus is depicted in Figure 2. The cellular stress marker ATF4 was significantly higher in the FGF23 treated group. A marker for ER-stress, namely the ratio of spliced over unspliced XBP1, was not significantly higher in the FGF23 group. However, for the s/uXBP1 ratio the p-value is highly influenced by one vehicle treated rat, this rat was in presence of the animals which showed behavioral responses, which possibly confounds stress measures. When excluding this rat, there is a significant difference between the FGF23 treated rats and the control group (p = 0.03). The other tested genes were not up- or downregulated after an icv FGF23 injection (i.e. *Npy*, *Pomc*, *Klotho*, *Fgfr1*, *Slc20a1* and *Slc20a2*). Exclusion of above-mentioned vehicle treated rat did not affect any of the other analyses. No mRNA expression was found for *Slc34a1* and *Slc34a2* in the arcuate nucleus.

**Figure 2.**
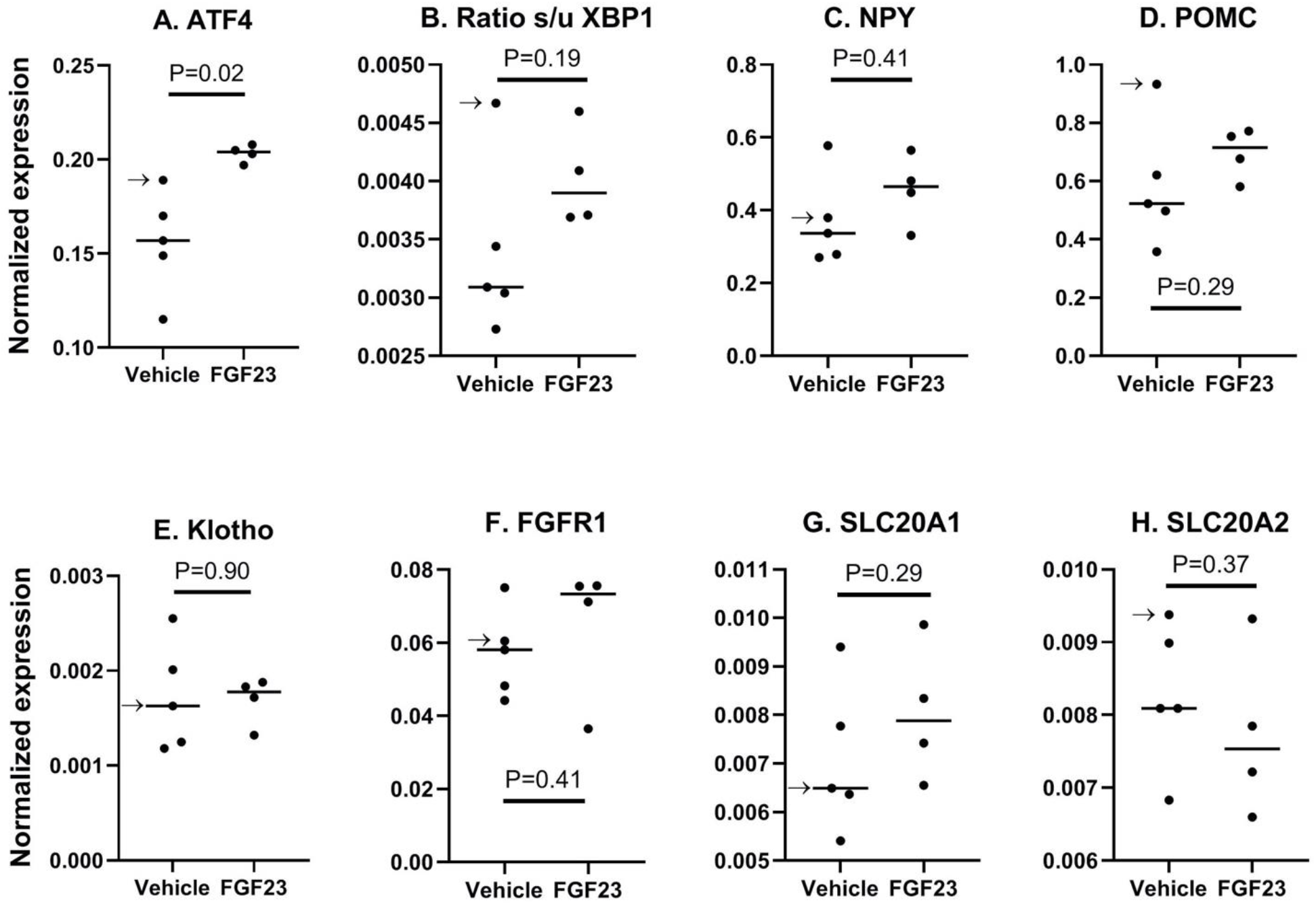
Normalized mRNA expression in the <u>arcuate nucleus</u> for (A) ATF4, (B) ratio of spliced/unspliced XBP1, (C) NPY, (D) POMC, (E) αKlotho, (F) FGFR1, (G) SLC20A1 and (H) SLC20A2 in FGF23 (n = 4) and vehicle (n = 5) treated rats. Bars reflect median values; the arrow indicates the rat which was present when higher doses of FGF23 was infused in rats and behavioral responses were observed.

In the lateral hypothalamus, ATF4 expression and the s/uXBP1 ratio did not differ between the tested groups (Figure 3). *αKlotho*, *Fgfr1*, *Slc20a1* and *Slc20a2* gene expression also showed no difference between the two groups. No mRNA expression was detected for *Slc34a1* and *Slc34a2*.

**Figure 3.**
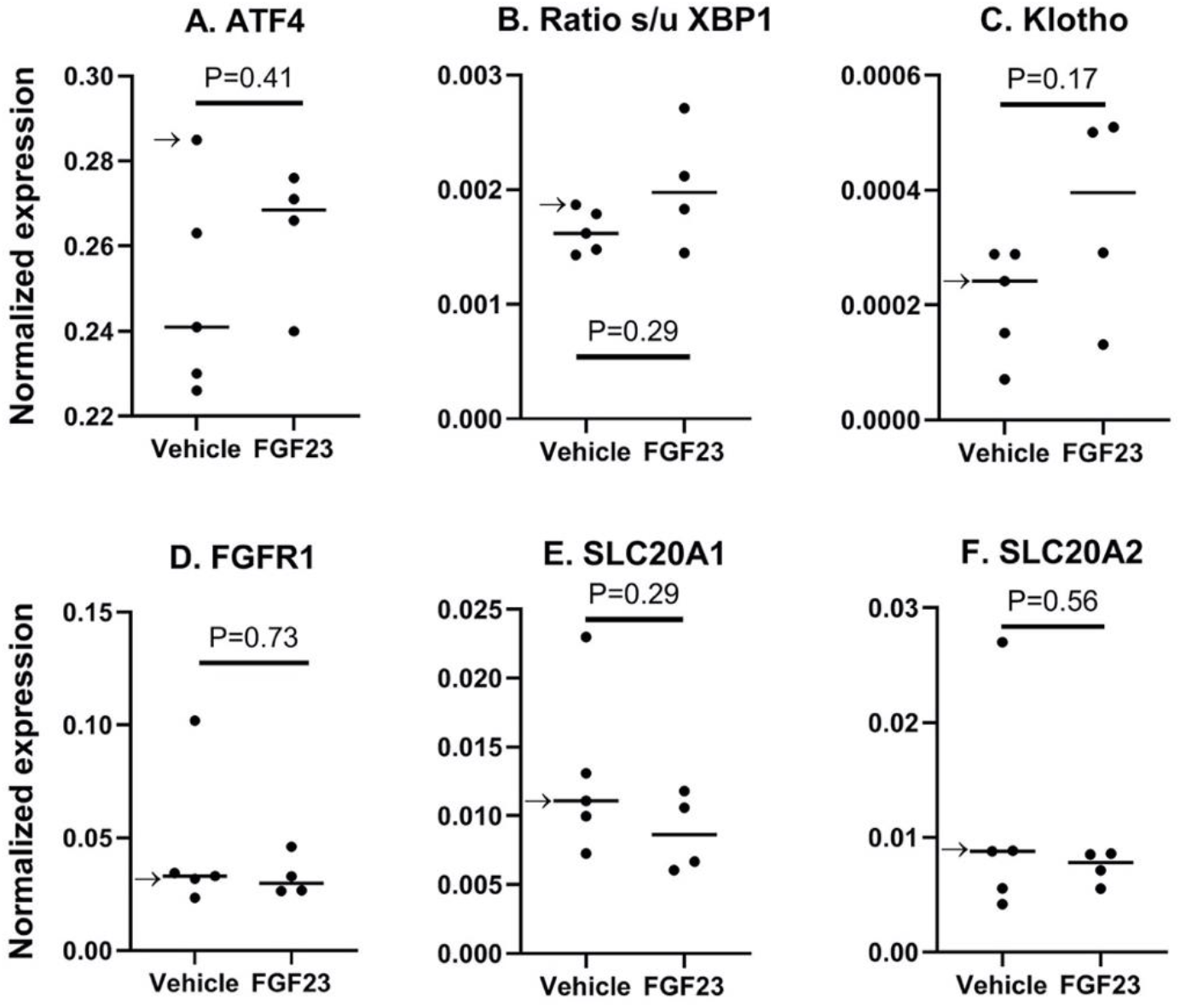
Normalized mRNA expression in the <u>lateral hypothalamus</u> for (A) ATF4, (B) ratio of spliced/unspliced XBP1, (C) αKlotho, (D) FGFR1, (E) SLC20A1 and (F) SLC20A2 in FGF23 (n = 4) and vehicle (n = 5) treated rats. Bars reflect median values; the arrow indicates the rat which was present when higher doses of FGF23 was infused in rats and behavioral responses were observed.

Gene expression in the choroid plexus is depicted in Figure 4. After FGF23 injection none of the tested genes differed in their expression compared to vehicle. In the choroid plexus *Slc34a1* mRNA was not detected.

**Figure 4.**
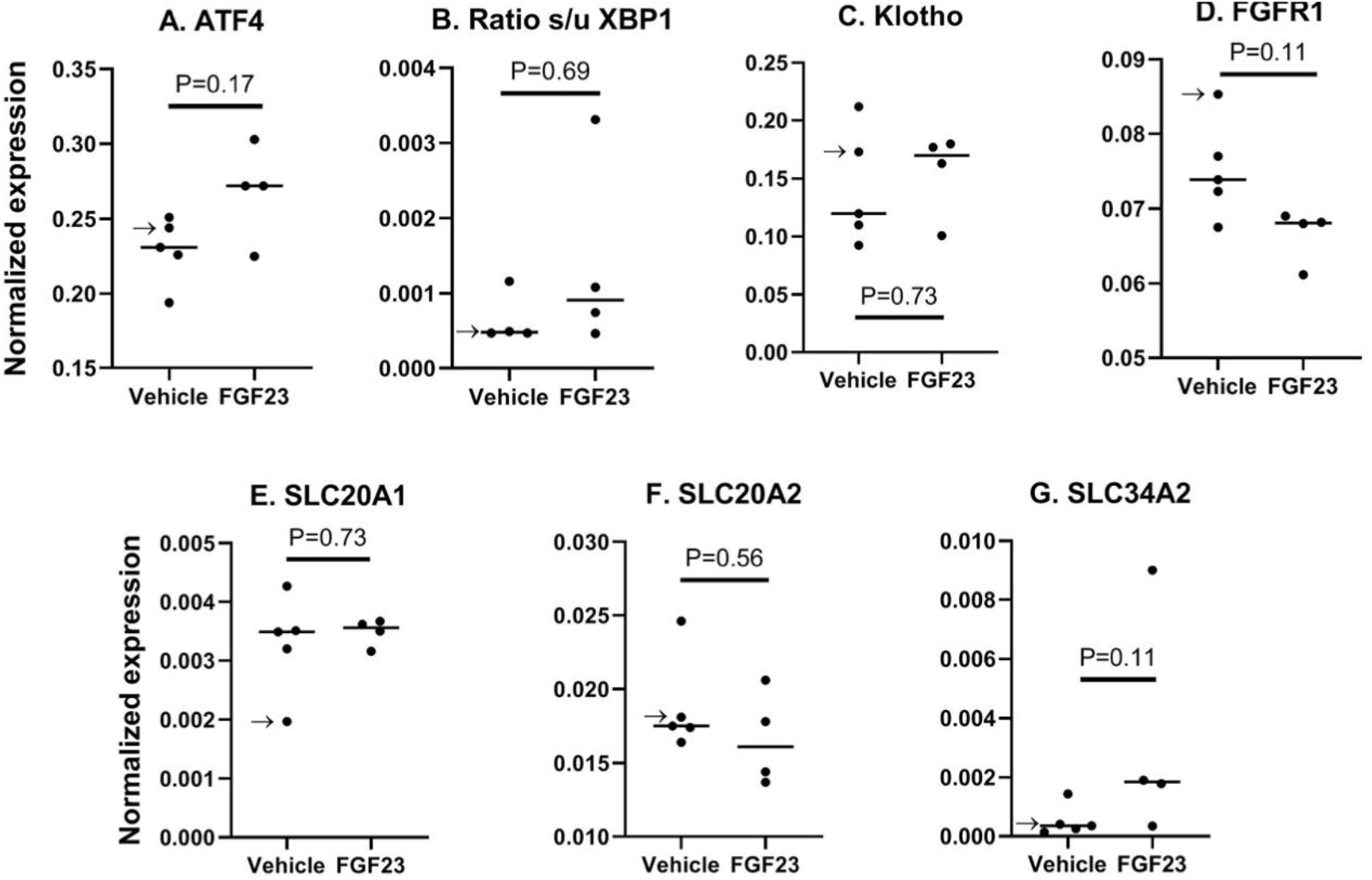
Normalized mRNA expression in the <u>choroid plexus</u> for (A) ATF4, (B) ratio of spliced/unspliced XBP1, (C) αKlotho, (D) FGFR1, (E) SLC20A1, (F) SLC20A2 and (G) SLC34A2 in FGF23 (n = 4) and vehicle (n = 5) treated rats. Bars reflect median values; the arrow indicates the rat which was present when higher doses of FGF23 was infused in rats and behavioral responses were observed.

## 4. Discussion

The objective of this study was to investigate the potential effect of FGF23 in the brain, with a first focus on NPY and POMC expression in the hypothalamus. To that end, FGF23 was administered in the third ventricle of rats and subsequent gene expression was studied in areas where FGF23 protein was identified earlier, being the arcuate nucleus, the lateral hypothalamus and the choroid plexus. Using a higher dose of FGF23, we observed clear behavioral effects that could be indicative of stress. Moreover, even with a lower dose, not exerting behavioral responses, we observed effects on markers of cellular stress at the side closest to the ventricle.

The behavioral changes and increases in gene expression of ATF4 and s/uXBP are likely due to effects of FGF23 in the brain and not because of other (chemical) substances present in the injectant. We had to use glycerol as a dissolvent to stabilize recombinant FGF23, which is a simple polyol, naturally present in the bloodstream. The toxicity of glycerol is very low, and it is well tolerated after intravenous or oral administration.^31^ Although little is known about its effect after central icv infusion, one study in rats found that long term icv infusion reduced body weight and food intake.^32^ However, the amount used was 12 times higher than the amount needed in our experiment as stabilizer. In a separate experiment we showed that icv injection of glycerol 10% is well-tolerated, for the amount needed as stabilizer. In subsequent experiments vehicle-treated animals received 10% glycerol as well.

The endoplasmatic reticulum (ER) is highly sensitive to cellular stress.^33^ Two important pathways induced in ER stress are the PERK and IER1 pathway, of which activation is reflected by the upregulation of ATF4 and increased splicing of XBP1, respectively.^34^ We observed a significant increase of ATF4 mRNA in the arcuate nucleus compared to controls. One animal received vehicle while other animals were treated with high-dose FGF23 at the same time, eliciting a behavioral response, which could have induced stress as rats are sensitive to environmental disturbances.^35^ If excluding this animal, XBP1 also increased significantly in the arcuate nucleus of FGF23 treated rats. Exclusion of this animal in all other analyses did not have a significant effect on the data. Further away from the injection site, in the lateral hypothalamus, no effect was observed on u/sXBP1 and ATF4. This was also the case for the choroid plexus. Taken together, these changes in cellular stress markers could point to an overall stress response.^36^ However, NPY and POMC mRNA were unchanged following FGF23 infusion and these are well known to be upregulated following psychological stress, rendering this hypothesis less likely.^37,38^

In addition, the absent effect on NPY and POMC are in contrast with preliminary findings in an earlier report.^22^ Morikawa *et al*. showed that after a 48h fast in mice, FGF23 protein increased in the hypothalamus and in serum.^22^ In a subsequent experiment, they administered icv FGF23 in 48h fasted mice. After 60 min, FGF23-induced ERK1 and AgRP immunoreactivity colocalized in the hypothalamus. At the same time, they observed an increase in NPY and POMC mRNA in the hypothalamus. Because of the extensive fast, these results must be interpreted with caution. Unfortunately, the data are derived from an abstract and the data were never published and thus details on methodology and dosing are not available.

In the kidney, FGF23 induces internalization and downregulation of two sodium-dependent phosphate channels: NaPi2a and NaPi2c. These are encoded by the SLC34A1 and SLC34A3 genes, respectively. As this is the main function of FGF23, it was previously hypothesized that FGF23 could also regulate phosphate homeostasis in the brain.^39^ Another important phosphate transporter family is encoded by SLC20A1 and SLC20A2 genes, which translate into sodium-dependent phosphate transporter 1 and 2 (PiT1 & PiT2). The SLC20 family is expressed in almost all tissues, including the brain, and is mostly viewed as a “housekeeping” transport protein.^40,41^ The SLC34 family is hardly detected in brain tissue, although one study found an unspecified NaPi2 transporter in the third ventricle.^41–43^ Phosphate is essential for normal cellular metabolic activity and its concentration in CSF is around 25% of the serum concentration.^44^ This gradient was suggested to be regulated by the SLC20 family in the choroid plexus.^44^ After administration of FGF23 in the third ventricle, we did not observe altered gene expression of the SLC20 family in the arcuate nucleus, lateral hypothalamus or choroid plexus. We did not detect gene expression of the SLC34 family in the lateral hypothalamus or the arcuate nucleus, although punches of the latter include cells of the third ventricle. In the choroid plexus we detected SLC34A2 mRNA, which was not affected by FGF23 administration. It should be noted that we only studied gene expression and can therefore not exclude that FGF23 had an effect on phosphate transporters on the protein level, such as internalization from the cellular membrane. This could be investigated in future research.

The behavioral effects we observed occurred fast and seemed to be transient. An explanation for this behavior could lie in acute effects on depolarization. Calcium has an important role in the CNS in regulating ion channel activity, neurotransmitter release, and depolarizing currents.^45^ In cardiomyocytes, FGF23 causes an acute peak in intracellular calcium concentrations *in vitro*.^46^ The effect of FGF23 on neuronal depolarization remains to be elucidated, however a recent study did observe FGF23’s depolarizing potential in neurons.^47^ In newborn Wistar rats, aged 0-5 days, the effect of FGF23 on the membrane potential of neurons in the rostral ventrolateral medulla (RVLM) was studied *in vitro*, in order to elucidate a possible mechanism behind the co-occurrence of hypertension and increased FGF23 levels in CKD.^47^ Indeed, FGF23 depolarized RVLM cells, whereas a FGFR1 antagonist caused hyperpolarization. In light of these findings, it could be that our observed effects upon icv FGF23 infusion *in vivo* indeed are caused by depolarization of specific neuronal populations, but this would warrant further research.

A limitation of this study is not being able to report on the effect of FGF23 in the brain on the protein level. As our primary goal was to determine effects on gene expression, tissue was used for mRNA extraction and no material was available for protein quantification. Hence, we were unable to look at effects of FGF23 infusion on several proteins. In addition, we could not isolate the third ventricle lining for further analysis, especially with regard to phosphate transporters. However, in punches of the arcuate nucleus, also the bottom of the third ventricle is included. In these punches, no effect was seen on gene expression of the phosphate transporters.

The main goal of the current study was to determine whether FGF23 affected gene expression within the hypothalamus involved in energy balance. This study has shown that a high dose of FGF23 had acute behavioral effects, suggestive for a stress response. Gene expression showed increased ER stress in the arcuate nucleus, but not in the lateral hypothalamus or choroid plexus. There was no effect on gene expression for the FGF23 receptor complex or phosphate transporters in all studied brain areas, also no effect was observed on NPY and POMC expression in the arcuate nucleus. In addition, we showed that use of glycerol as stabilizer for icv injections is well-tolerated. A natural progression of this work is to analyze the effect of icv FGF23 injections on hypothalamic protein levels, as well as systemic glucose tolerance and food intake.

